# Wall-associated Receptor Kinase and The Expression Profiles in Wheat Responding to Fungal Stress

**DOI:** 10.1101/2021.07.11.451968

**Authors:** Hong Zhang, Hanping Li, Xiangyu Zhang, Wenqian Yan, Pingchuan Deng, Yining Zhang, Shanlin Peng, Yajuan Wang, Changyou Wang, Wanquan Ji

## Abstract

Cell wall-associated kinases (WAKs), which are encoded by conserved gene families in plants, are crucial for development and responses to diverse stresses. However, the wheat (*Triticum aestivum* L.) WAKs have not been systematically classified, especially those involved in protecting plants from disease. Here, we classified 129 WAK proteins (encoded by 232 genes) and 75 WAK-Like proteins (WAKLs; encoded by 109 genes) into four groups, via a phylogenetic analysis. An examination of protein sequence alignment revealed diversity in the GUB-domain of WAKs structural organization, but it was usually characterized by a PYPFG motif followed by CxGxGCC motifs, while the EGF-domain was usually initiated with a YAC motif, and eight cysteine residues were spliced by GNPY motif. The expression profiles of WAK-encoding homologous genes varied in response to *Blumeria graminis* f. sp. *tritici* (*Bgt*), *Puccinia striiformis* f. sp. *tritici* (*Pst*) and *Puccinia triticina* (*Pt*) stress. A quantitative real-time polymerase chain reaction (qRT-PCR) analysis proved that *TaWAK75* and *TaWAK76b* were involved in wheat resistance to *Bgt*. This study revealed the structure of the WAK-encoding genes in wheat, which may be useful for future functional elucidation of wheat WAKs responses to fungal infections.

## Introduction

To survive and reproduce, plants have evolved a cassettes of adaptive mechanisms in response to stresses from environmental changes. Cell wall (CW) is a paramount plant’s structure that fences and protects the plant cell, and it has also emerged as a component of important systems that monitor plant growth and defense [1, 2]. A class of important proteins residing in plasma membrane is receptor-like kinases (RLKs) or receptor-like proteins (RLPs), which are developed for maintaining the cell wall integrity (CWI) [3, 4]. Cell wall-associated kinases (WAKs), a subgroup of RLKs, link the plasma membrane to the carbohydrate matrix [5]. Another subgroup of RLKs is the key factors of pathogen defense mechanisms that were collectively referring to as pattern recognition receptors (PRRs)[6, 7]. Conceptually, apart from microbial effectors, PRR proteins recognize pathogen associated molecular patterns (PAMPs) that are conserved motifs from pathogens [6] and ‘plant-self’ derived damage associated molecular patterns (DAMPs) during pathogen development [1, 8]. Generally, RLKs contain an N-terminal signal peptide, a variable extracellular domain (ECD), a single-pass transmembrane domain, and a cytosolic protein kinase domain [9]. WAKs contain an extracellular GUB-WAK binding domain and an EGF (Epidermal Growth Factor) domain interacting with pectin [10, 11] and oligogalacturonides (OGs) [8, 12], while PRRs characterized with LRR domain in non-cytoplasmic N-terminal region[13, 14]. Intriguingly, both WAKs and PRRs trigger directly signal cellular events through their cytoplasmic kinase domain, which hinted that plant defense mechanism may be overlapped or have a crosstalk between WAKs with PRRs. Recently, although the WAKs are generally required for cell expansion, the evidence that WAK genes participate to resistance to pathogens in many plant species against fungi and bacteria had been well provided [15–19]. Plant pathogens secrete a battery of cell wall-degrading enzymes (CWDEs) to degrade cell wall for infection [20], whereas, such host-derived cell wall degradation products can become DAMPs and then elicit plant immune responses [21]. These acknowledge guaranteed the further study on the role of WAKs in plant disease defense, including whether WAKs associate with other RLKs to ensure appropriate function, especially with PRRs.

The WAK genes belong to large gene subfamilies. Twenty-six *AtWAKs* and *AtWAKLs* in *Arabidopsis* [22], 125 *OsWAKs* in rice (*Oryza sativa*) [23], 91 WAKs in barley [24], 44 members in Apple [25] and 16 putative *GhWAKs* from *Gossypium hirsutum* [26] were identified based on structural domain identification. The expression of WAKL5 and WAKL7 can be induced by salicylic acid analog in an nonexpressor of pathogenesis-related gene 1 (NPR1)-dependent manner, suggesting that they are PR genes [27]. *AtWAK2* can bind pectin and activates the mitogen-activated protein kinases (MPK) in Arabidopsis [28]. Overexpression of *OsWAKL21.2* activates immune responses in both rice and Arabidopsis [18]. Using rice loss-of-function mutants of three co-regulated genes *OsWAK14, OsWAK91* and *OsWAK92*, Cayrol et al. proved that individual OsWAKs are positively required for quantitative resistance to the rice blast fungus (*Magnaporthe oryzae*)[15]. A *ZmWAK* gene located in the loci of *qHSR1* confers quantitative resistance to maize head smut [29]. Wheat (*Triticum aestivum* L.) is one of the four major cereals in the world, whereas its growth and production are severely affected by pathogen [30], such as stripe rust (*Puccinia striiformis* f. sp. *Tritici, Pst*), powdery mildew (*Blumeria graminis* f. sp. *tritici; Bgt*), head blight (*Fusarium graminearum*), sheath blight (*Rhizoctonia cerealis*) and take-all (*Gaeumannomyces graminis*) [31]. Wheat is a hexaploid species with a large and complex genome, which means that it has more number of WAKs coupling more complex coordination between them. However, at present, only one WAK-like protein, WAKL4/Stb6 was substantiated to confer pathogen resistance to *Zymoseptoria tritici* without a hypersensitive response [19]. According to the cues given by other plant species, we could infer the WAKs that are involved in the elaboration of cell wall damage-induced immune responses remain poorly understood in wheat. Here, we advocated that the identification and understanding of association between WAKs, CWI sensors and trigger immunity (TI) related PRR-DAMP pairs in wheat will help us to designate agricultural strategies through developing wheat varieties with enhanced disease resistance.

## 2. MATERIALS AND METHODS

### 2.1 Plant materials and pathogen stress treatment

The N9134 winter wheat cultivar, which is resistant to all *Bgt* races in China, was crossed seven times with a susceptible recurrent parent, Yuanfeng (YF175), to obtain a resistant line with the YF175 background. This new resistant line was named YF175/8*PmAS846. To further eliminate the influence of inhomogeneity, a BC_8_F_1_ resistant plant was self-crossed to generate the contrasting BC_8_F_2_ homozygous lines, which differ only regarding *PmAS846* on chromosome 5BL, and named N9134R_1_ and N9134S_1_. Additionally, *Bgt* isolate E09 was maintained on SY225 wheat plants. The contrasting *Bgt* RILs N9134R/S and SY225 plants were cultivated in soil in a growth chamber at 18 °C with a 16-h light/8-h dark photoperiod. Seedlings at the 3-leaf stage were inoculated with *Bgt* conidia from SY225 seedlings that were inoculated 10 days earlier. The *Bgt*-inoculated N9134R/S leaves were harvested at 6, 12, 24, 36, 48, 72, and 96 hours post-inoculation (hpi), after which they were immediately frozen in liquid nitrogen and stored at −80 °C. Subsequent analyses were completed with three biological replicates. For the genome-wide transcription analysis, RNA-Seq Data generated from the inoculated N9134 leaves were harvested at 0, 24, 48, and 72 hpi as previously described [32]. The non-inoculated sample (0 hpi) was used as the control.

### 2.2 Identification and sequence analyses of wheat WAK proteins

To obtain detailed information regarding wheat WAKs, we downloaded all available sequences for proteins annotated with ‘WAK’ in the EnsemblPlants database (http://plants.ensembl.org/Triticum_aestivum/Info/Index). The Arabidopsis WAK protein sequences were also used as the queries for a BLASTp search against Chinese Spring protein databases and NCBI database. After screening EBI database (http://www.cbs.dtu.dk/services/) with the InterProScan [33], TMHMM and SignalP, all downloaded protein sequences were analyzed using Conserved Domain Search Service (CD-Search) software in NCBI (https://www.ncbi.nlm.nih.gov/Structure/cdd/wrpsb.cgi). Furthermore, the candidate proteins were identified based the condition of signal peptides (SP) in N terminal, cysteine-rich galacturonan-binding domain (GUB-WAK, pfam13947 and/or cl16494), EGF-like domain (EGF_CA, cd00054), TMhelix and following protein kinase or serine/threonine protein kinase (PKC/STKc, cl21453 or cd14066) domains. Furthermore, all of the remaining sequences was screened in EnsemblPlants database with BLASTP (Protein Basic Local Alignment Search Tool) algorithm for sequence matches (85% identity as the threshold). And then the WAK proteins were sorted and classified after redundant sequences were manually removed. The motifs of the encoded WAK proteins were detected using the motif-based sequence analysis program (MEME) [34], whereas a phylogenetic neighbour-joining tree was constructed for the HSP proteins with the molecular evolutionary genetics analysis (MEGA) program (version 6.0) [35].

### 2.3 WAK genes expression analysis

To detect the expression of *WAKs* responding to fungal stress, the transcriptome data of 21 samples (the leaves of N9134 infected by *Pst* and *Bgt* at 0, 24, 48 and 72 hpi, Accession No. PRJNA243835 in NCBI) were analyzed [32], while the 15 sets of data of resistance line Avocet*Yr5 inoculated with *Pst* (at 0, 24, 48, 72 and 120 hpi) were obtained from the NCBI database with accession No. PRJEB12497 [36]. Additionally, 15 sets of data of resistance line WL711+Lr57 inoculated with leaf rust pathogen (*Puccinia triticina, Pt*) (at 0, 12, 24, 48, and 72 hpi) were also downloaded from the NCBI database with accession No. PRJNA328385 [37]. The expression level of WAK-encoding genes that involved in wheat responses to fungal infections were screened using the reads per kilobase of exon model per million of aligned readings (RPKM) values for each transcription in sample. In addition, the expression profiles of the specific *TaWAKs* in the infected leaves of the contrasting NILs were analysed by a SYBR Green-based qRT-PCR with cDNA prepared from samples collected at 6, 12, 24, 36, 48, 72, and 96 hpi. The uninoculated plant samples at the same time points were set as the controls. Three independent biological replicates were prepared for each time point. Primers specific for the examined WAK-encoding genes (Supplemental **Table S1**) and *TaActin* (i.e., internal reference) were designed with the Primer 5 program, and were used to analyze gene expression levels. The qRT-PCR was completed with the FastKing RT Kit (with gDNase) (TIANGEN, Beijing) and the QuantStudio™ 7 Flex Real-Time PCR System (Life Technologies Corporation, USA). The PCR reaction and program were carried out as previously reported. For each sample, reactions were completed in triplicate, with three non-template negative controls. A standard 2^−ΔΔCT^ method was used to quantify relative gene expression levels.

## 3. RESULTS

### 3.1 Identification of wheat WAK family proteins

In order to elucidate the mechanism mediating wheat resistance to *Bgt*, we recently predicted the key genes based on a WGCNA, which contained 114 nodes proteins. Notably, cell wall-associated kinase was annotated as a markedly node protein in this net [38] with the connectivity value reaching to 1583.8. Screening the PmAs846 physical map (from 541528564 to 546124256) with the Chinese Spring reference genome, indicated that 7 out of 57 genes were annotated wall-associated receptor kinase or pseudogene of WAKs between the co-segregation markers BJ261635 with XFCP1 [39]. To further evaluate the genomic distribution and function of WAKs in wheat responding to fungi, here we investigated WAK-encoding genes in genome-wide (**Fig. S1**). The protein sequence homology resulting data confirmed the presence of 204 encoding proteins with a characteristic GUB-WAK binding domain in the wheat genome, which usually accompanied with C-terminal PKC or STKc domain. Of which, 129 proteins (encoded by 232 genes) contain EGF domains as well as SP, GUB-WAK, internal TMhelix, PKC/STKc. Therefore, according to the protein structure, these 129 proteins are designated as TaWAKs, while the others 75 proteins (encoded by 109 genes) absenting EGF-CA, SP and/or TMhelix are classed into TaWAKLs. These 204 proteins were encoded by 341 wheat genes and distributed in the terminal of 20 chromosomes (**Figure 1**). Specific details are provided in Supplemental **Table S2**. TaWAKs were named considering the orthology to the reported WAKs and homoeology between A-, B- and D-subgenomes. Since only a few TaWAKs were reported, the most of names were given according to the order on the seven partial homologous group. Lower case letters (a, b and d) is employed to differentiate homologous genes from A-, B- and D-subgenome. The additional number (1 to 4) is added after the letter to indicate the repetition of the WAK protein-encoded genes on the same chromosome.

**Figure 1.**
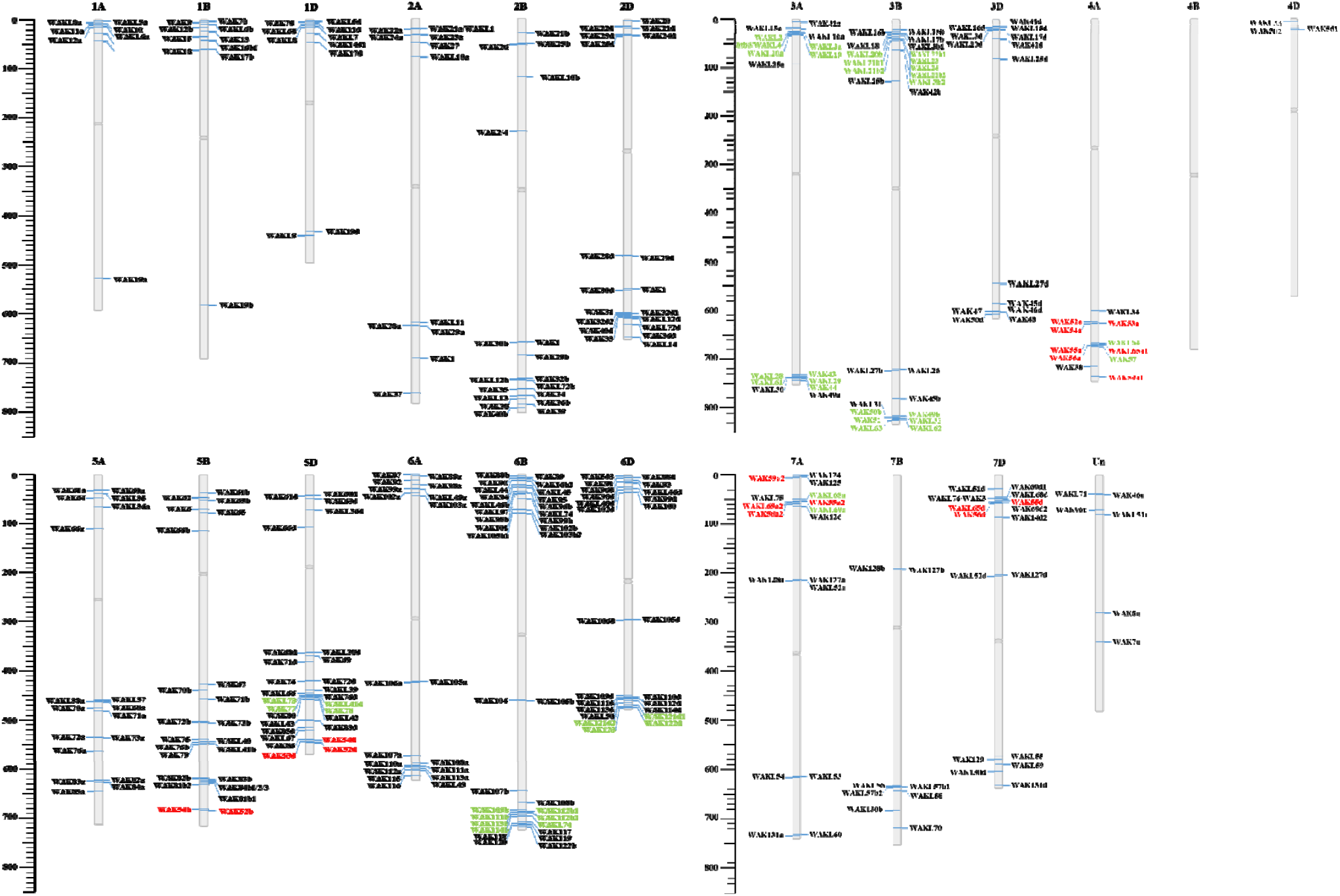
Chromosomal locations of WAK genes in wheat. A total of 341 *TaWAKs* were localized to *Triticum aestivum* L. The homologues genes that resided in differential homologues group were marked with red characters. WAK genes in tandemly arrayed gene (TAG) cluster were marked with green characters.

#### Phylogenetic analysis and evolution of WAK family proteins in wheat

To elucidate the evolutionary relationship of WAK family proteins in wheat, we performed a phylogenetic analysis using the tool MEGA X[40]. The results showed that wheat WAKs were interleave with the isolated clade formed by the five Arabidopsis AtWAKs and ten cotton GhWAKs (**Figure S2**). This suggests that some members of WAKs in wheat have likely evolved more ancestral than that in Arabidopsis, and subsequently another part of members have evolved with AtWAKs together independently. Based on the phylogenetic relationships, we divided the wheat WAK members into three classes (I to □) according to the isolated main clade from Arabidopsis and/or cotton. Among these TaWAKs, the most of wheat WAKs could be classified in two clade I and □, with TaWAK1, TaWAK30, and TaWAK65 in clade □; TaWAK71, TaWAK109, TaWAK110, TaWAK111, TaWAK112, TaWAK113, TaWAK114, and TaWAK115 in clade □ (**Figure S2**). Collectively, these members of clade I and □ have evolved in different trajectory form that in clade □ and □.

Screening WAK genes showed that WAKs distribute unevenly among chromosomes as tandemly arrayed gene (TAG) cluster in *Arabidopsis* and cotton. Here, we investigated the physical positions of *TaWAK* genes and found that *TaWAKs* was also unevenly distributed in seven homologous groups with some members as TAG clusters (more than 3 consecutive WAKs). A significant characterization is that nearly all of TaWAK-encoding genes are distributed at the distal of chromosome (**Figure 1**). A total of nine WAK-TAG clusters presented on chromosome 3A, 3B, 4A, 5D, 6B, 6D and 7A. Of which, there have four WAK-TAGs in chromosome 3A and 3B, while no WAK-TAG was detected in group 1 and 2. The number of *TaWAKs* reach to 77 in the homologous group 6, while only 14 were identified in the group 4. Notably, 11 of 14 *TaWAKs* in group 4 were encoded by the genes from chromosome 4A, and no any *TaWAKs* was identified on the B subgenome (i.e. 4B). Intriguingly, three WAKs on 4A (*TaWAK52a*, *TaWAK53a* and *TaWAK54a*) are high homologue to the genes of chromosome 5B/D, while 5 WAKs are homologue to that of 7A/D (**Table S2** and **Figure S3**). This suggested that the fragment of chromosome 4AL was substituted with a partial homolog fragment of 5AL and 7BS during speciation and/or evolution of hexaploid *T. aestivum*. Additionally, it is commonly that the TAG clusters were composed of *TaWAKs* and *TaWAKLs* together, for example: *TaWAKL64*, *TaWAK55*, *TaWAKL65*, *TaWAK56* and *TaWAK57* presenting as a cluster of TAGs on chromosome 4A; *TaWAK49*, *TaWAK50b*, *TaWAKL32*, *TaWAK51*, *TaWAKL62* and *TaWAKL63* clustering in 3B.

Furthermore, we analyzed the protein domain structures of TaWAKs (Supplemental **Figure S4**). Alignments of the amino acid frequencies of the EGF_like domains in 129 TaWAKs revealed ten highly conserved cysteine (Cys) residues (YACxxxxxxCxxxxxxxGYxCxCxxxYxGN[PA]Y[LI]xxGCx[DN][IV][DN]ECx_4-7_Cx_1-3_GxCx[ND]xx GxxxCxCxxGxxGx, where x is any amino acid) (**Figure 2**), which are implicated in the formation of disulfide bonds and/or required for calcium binding. The WAK domain is the most variable of all the domains contained within a WAK protein; however, our results indicated several conserved residues near the N- and C-terminal ends of WAK domain (N-terminus: xCxxxCGxxx[IV]PYPFGx_4-9_[CS]xxxxFxxxCxxxxxxPx; C-terminus: Nx[LF]xxxG Cx_24-42_GxCxxxGCC) of this domain in TaWAK proteins (**Figure 2**), which share a high level of similarity with that in AtWAKs and GhWAKs [26].

**Figure 2.**
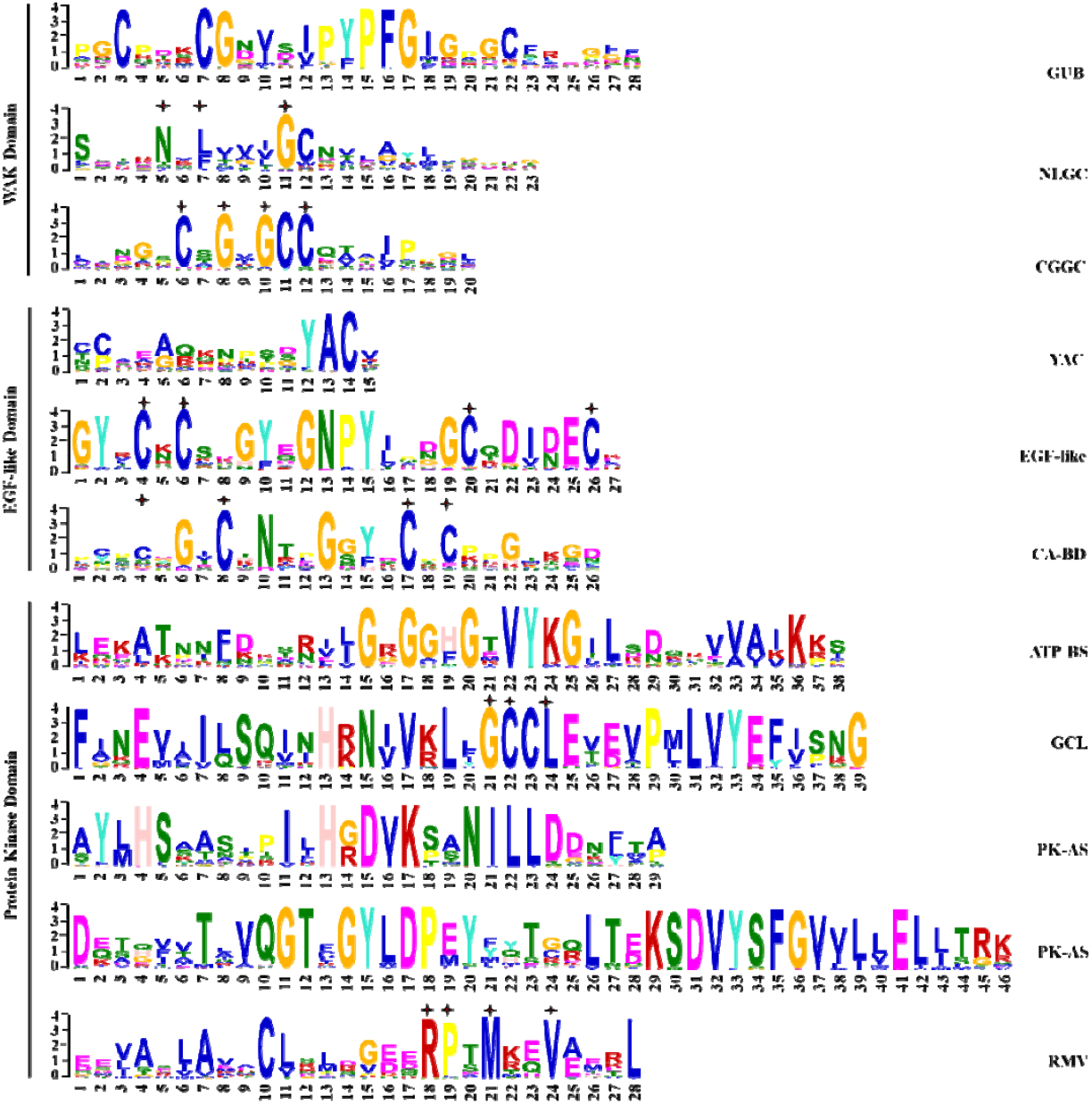
The conserved functional domains and motifs in WAK proteins of wheat. The characterization of amino acids was identified by multiple alignment using MEME software. The conserved amino acids of motifs were marked with plus signal above the letters. The name of motifs was indicated at the right of each sequence.

#### Expression of TaWAK protein-coding genes involved in wheat resistance against fungal pathogens

To identify the WAK-encoding genes involved in wheat responses to *Bgt*, *Pst* and *Pt* infections, we screened 60 RNA-seq libraries (PRJNA243835, PRJEB12497 and PRJNA328385). Based on the RNA-seq data of wheat under pathogen infection, the expression heatmap of *TaWAKs* on chromosomes A, B and D were made by TBtools after simply eliminating the non-expressed genes. Under the stress of *Bgt*, 28 *TaWAKs* expression were significant dysregulated after infection (**Figure 3**A), while 23 and 13 *TaWAKs* were identified as DEGs after *Pst* and *Pt* infection, respectively (**Figure 3B, 3C** and Supplemental **Table S3**). Notably, 3 *DE-WAKs* were overlapped in three pathogen species infection (**Figure 3**D), including *TaWAK32b, TaWAK69* and *TaWAK70a*. This means that the three *WAKs* may be involved into basal resistance to fungi.

**Figure 3.**
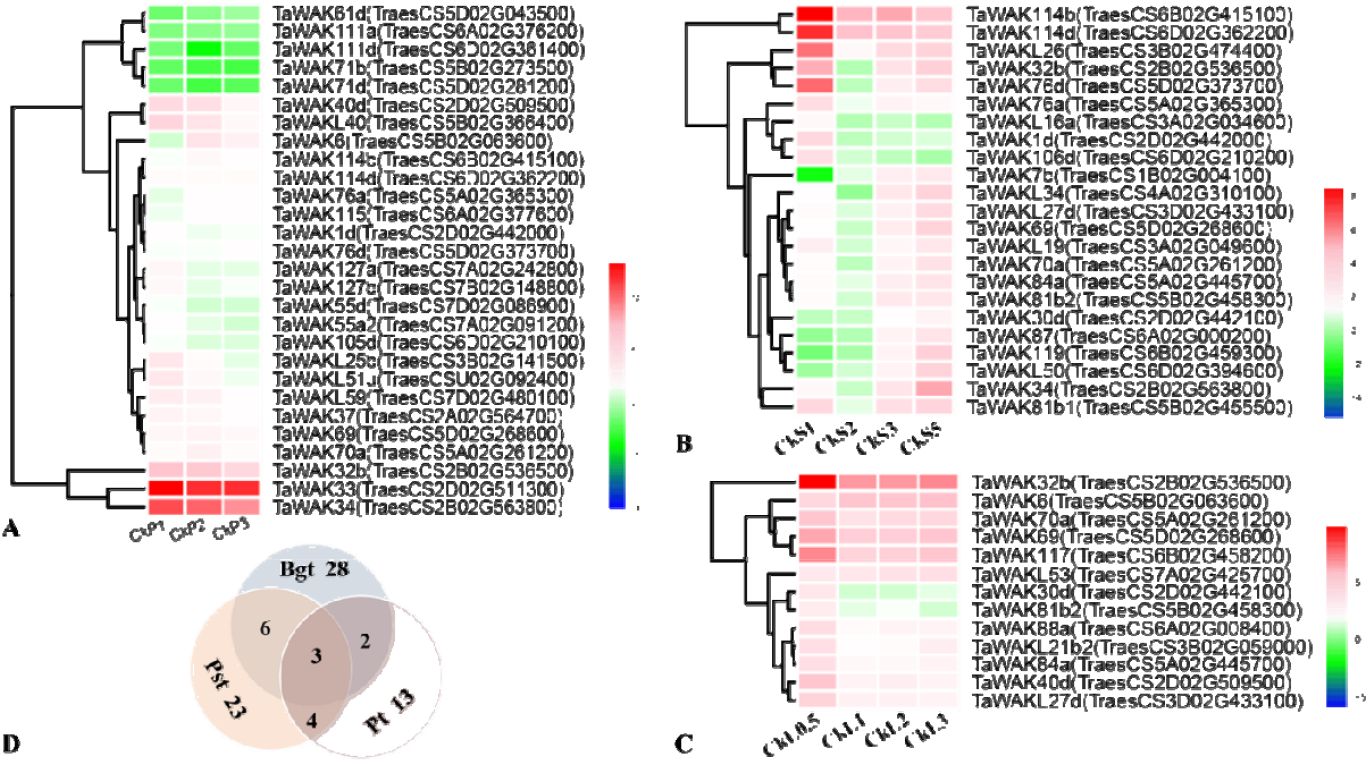
Heat map of gene expression of *DE-TaWAKs*. A: N9134 responds to Bgt infection at 24, 48 and 72 hpi. B: Avocet*Yr5 responds to *Pst* infection at 24, 48, 72 and 120 hpi. C: A resistance line WL711+Lr57 responds to *Pt* infection at 12, 24, 48 and 72 hpi. D: Venn diagram of different fungi pathogen infection on resistance line. The expression change was presented with log2(FC) at corresponding time points comparing to that at 0 hpi. Gene No were listed at the right of the corresponding expression, while the bar scale was given to represent the value of log2(FC).

Of which, the expressions of *TaWAK76a* and *TaWAK76d* are always up-regulated 2.2 to 3.6 times during infection stage (at 24, 48 and 72 hpi) than that in control (**Figure 3A**). Notably, *TaWAK76a* (TraesCS5A02G365300.1) and *TaWAK76d* (TraesCS5D02G373700.1) are homolog genes to *TaWAK76b* (TraesCS5B02G366900.1), residing in the genetic fragment of PmAS846 (carried by N9134 and flanked by STS markers *BJ261635* and *Xfcp620*, i.e. from TraesCS5B02G361900 to Traes5B02G368200). Additionally, another WAK gene, *TaWAK75* (TraesCS5B02G363100.1) was missed in N9134 due to the substitution with a fragment of Wild emmer AS846 so that its’ expression could not be detected using CS reference genome. Considering that the similarity of expression trends of subgenome homologous genes was observed in wheat after fungal infections [41], we examined the expression of *TaWAK75* and *TaWAK76b* in N9134R_1_ and N9134S_1_ (YF175/7*PmAS846, a YF175 backcross line carrying PmAS846 from N9134 or not).

The expression profiles of *TaWAK75* and *TaWAK76b* in the inoculated N9134R_1_ and N9134S_1_ plants are presented in **Figure 4**. Following the inoculation with *Bgt*, the expression pattern of *TaWAK76b* exhibited both in the on-and-off manner. The expression levels of *TaWAK76b* were significantly supressed at 6 hpi in line N9134R_1_ and N9134S1. However, the double peaks were detected at 12 and 96 hpi in resistance background, while the accumulated transcription level in susceptible line at 24 and 48 hpi reach to the maximum, which are 26 and 45 times higher than that in uninfected control respectively. For the expression of *TaWAK75*, the transcription level was stably supressed at 6, 12, 24, 36, 48, 72 and 96 hpi after *Bgt* inoculation in N9134S1. At 12 hpi, the related expression quantity touch to the bottom, which was 50 times lower than that in control. This means that *TaWAK75* and *TaWAK76b* were involved into wheat responding to *Bgt* infection. The former maybe play a negative effector role here, while the latter functioned in the fundamental resistance upon the pathogen successfully punctured into plant cell.

**Figure 4.**
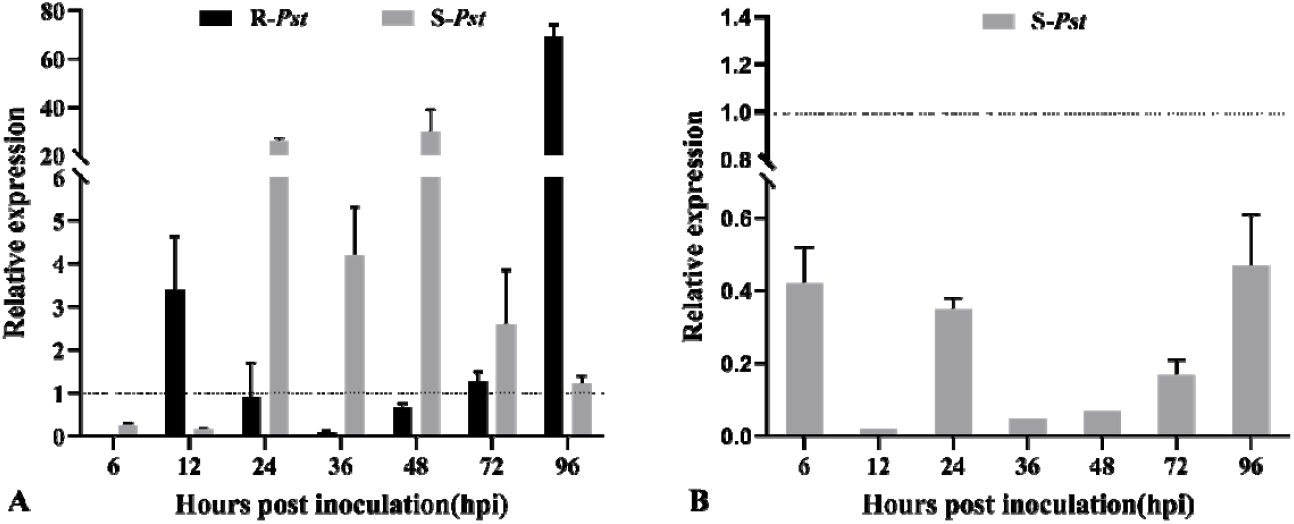
Expression patterns of *TaWAK75* and *TaWAK76b* in N9134R_1_ and N9134S_1_ induction by powdery mildew pathogen E09. Gene expression levels were assessed by qRT-PCR at 6, 12, 24, 36, 48, 72, and 96 hpi. Data were normalized to the actin expression level. The mean expression value was calculated from three independent replicates, and the standard deviation was given at each time points. A represents *TaWAK76b* gene expression in N9134R_1_ and N9134S_1_ after powdery mildew E09 infection; B represents gene expression of *TaWAK75* in susceptible line N9134S_1_. The dotted line indicated the controls at the same time points. The names of corresponding gene were listed in the top of each panel.

## Discussion

WAKs are primarily involved in regulating cell expansion and protecting plants from detrimental effects depending upon the state of pectin [10], and also involved in the response to pathogens. For example, induction of *AtWAK1* is required for plants to survive *Pseudomonas syringae* infection [42], while the *OsWAK1* transcripts was significantly induced by *Magnaporthe oryzae* [43]. *AtWAKL10* is co-expressed with pathogen defense related genes and is strong and rapid induced by a range of pathogens and their characteristic MAMPs including chitin and flg22 [44]. CsWAKL08 confers resistance to citrus bacterial canker via ROS control and induced by phytohormones salicylic acid (SA) and methyl jasmonic acid (MeJA) [45]. Overexpressing analysis of *TaWAK6* in plant-pathogen interactions showed a lower number of uredinia and higher rates of necrosis at the *Pt*-infected sites [46]. These indicated that *WAKs* may function in MAMP initiated basal defense responses to a broad range of pathogens in similar or different fashion. However, the detailed possible involvement in plant-fungal mutual interaction is yet to be established. Notably, PTI pathways activated by cell-surface and ETI intracellular receptors in plants mutually potentiate to activate strong defenses against pathogens [47, 48]. It is no doubt that understanding the role of WAKs in plant responding to fungal stress will helpful in disclose the mechanism of immunity. Here, we firstly identified WAK and WAKL family proteins (129 and 75 respectively) in common wheat, while the current study generated useful information regarding the expression of 46 WAK-encoding genes in responding to fungi infection. The results showed that the member of TaWAK proteins is nine times more than that in *Arabidopsis*, and double than that in *O. sativa*. In addition, three sub-genomes characterization of wheat destined to complex the function of WAK-encoding genes (*232 WAKs* and 109 *WAKLs*). The presented data may form the basis for future investigations into the *WAKs* expression dynamics during plant–pathogen interactions. Furthermore, our findings may be relevant for further elucidating the role of WAKs in wheat defense responses as well as the detailed effects of the diversity.

The extracellular EGF domain in WAK known to bind with high affinity to specific cell-surface receptors (i.e. small peptides) and form homo and heterodimers with receptors [49, 50]. In plant, PRR proteins recognize pathogen derived PAMPs and host-derived DAMPs during pathogen invasion, which induce PAMP triggered immunity (PTI) or basal defense [51]. It is noted that PRR/DAMP protein complex interact with AtWAK1 and OGs [8], a derivatives of pectin which is the methylated ester of polygalacturonic acid (PGA) acting as a cementing material in the cell walls of all plant tissues. Notably, fungal cell wall is mainly composed of polysaccharides (e.g. chitin, glucan) and proteins [52]. It is demonstrated that the transcriptional regulation of the WAK genes could be triggered by various active ‘effectors’ including exogenous and endogenous elicitors. Here, we found that 46 *TaWAKs* was significantly induced by *Bgt*, *Pst* and/or *Pt* invasion. Intriguingly, more than 30% *DE-WAKs* were overlapped in responding to three fungi infection (Fig. 3D), while *TaWAK32b* (TraesCS2B02G536500), *TaWAK69* (TraesCS5D02G268600) and *TaWAK70a* (TraesCS5A02G261200) were commonly induced by three fungi with similar expression pattern (Fig. 3A, 3B and 3C). Considering the very low expression level of them at 0 hpi (supplemental Table S3), we inferred that they should be fundamental RLKs to detect fungal hyphae invasion, although the further substantial experiment need to be tested. A common feature of all EGF-like domains is that they are found in the extracellular domain of membrane-bound proteins or in proteins known to be secreted. Many of membrane-bound and extracellular proteins require calcium for their biological function and a calcium-binding site has been found at the N terminus of some EGF-like domains [53]. Calcium-binding may be crucial for numerous protein-protein interactions [54]. The EGF-like domain includes six cysteine residues in human which have been shown to be involved in disulfide bonds. However, we found *WAKs* in wheat contained nine conserved cysteine residues following YAC motif at the N-terminal, which is similar with *GhWAKs* [26]. Notably, the YAC motif is totally absented from *AtWAKs*. The recently reports showed that GhWAK7A directly interacts with both GhLYK5 and GhCERK1 (lysin-motif-containing RLKs) promoting chitin-induced GhLYK5-GhCERK1 dimerization and phosphorylates GhLYK5 [26]; a *ZmWAK* confers quantitative resistance to head smut [29], while two WAK like genes, *XA4* and *Stb6*, conferred resistance to *Xanthomonas oryzae* pv. *oryzae* (*Xoo*) and *Zymoseptoria tritici*, respectively [19, 55]. These set important examples for the role of WAK in plant resistance to pathogen. In addition, our previously WGCNA analysis predicted that a WAK partial protein (TraesCS5B02G160500) was a subcentre node in *Bgt*-infected N9134, which coupling with RLP37, RPM1 and RPS2 [38]. Taken this information together, the results presented here demonstrate that *TaWAKs* may be an indicator that links plant perceiving or innate immune defense capacity in hexaploid wheat as well.

In conclusion, in this study, we reveal the immense complexity of *WAKs* underlying the responses of wheat to fungal infection. We identified 204 TaWAK proteins and performed transcriptome expression analysis using 42 sets of RNA-seq data. Although it is not clear why WAKs was intensely induced by powdery mildew infection in immune wheat leaves, and why the levels of expression were lower than in susceptible wheat, our results substantiated the possibility that WAKs has a broad spectrum of functions in plants. In conclusion, this insight into WAKs gene expression will be useful in dissecting the mechanism of fungal resistance in wheat, and understanding the interaction between cell wall and pathogen defense.

## SUPPLEMENTAL DATA

Supplemental Table S1. Primers used for the qRT-PCR analysis of WAK-encoding genes.

Supplemental Table S2. The list of wheat WAK proteins.

Supplemental Table S3. The expression level of differential expressed WAK-encoding genes.

Supplemental Figure S1. Genome-wide identification pipeline of WAKs in wheat genomes.

Supplemental Figure S2. Phylogenetic tree of WAK proteins from wheat, Arabidopsis and cotton.

Supplemental Figure S3. Phylogenetic tree of WAK homologues proteins from differently partial homologous group of wheat.

Supplemental Figure S4. Phylogenetic tree, conserved motifs, and sequence structure of 129 TaWAK proteins.

## Conflicts of Interest

The authors declare that they have no conflict of interest.

## Authors’ contributions

HZ and WJ conceived the project and provided overall supervision of the study; HZ, PH, WY, YZ and SP performed the experiments and data analysis; YW and CW contribution to developing the materials; XZ and PD helped in experimental works and analyzed the RNA-Seq data; HZ wrote the first version of the paper; all authors reviewed and approved the final manuscript.

## Acknowledgments

This work was financially supported by Natural Science Foundation of Shaanxi Province (grant no. 2021JM-090).

## Notes

### Competing Interest Statement

The authors have declared no competing interest.

