## supplemental file 1 for "Wall-associated Receptor Kinase and The Expression Profiles in Wheat Responding to Fungal Stress"

**Table S1.** Details regarding primers used for the qRT-PCR analysis of WAK-encoding genes.

| Primer | Forward primer sequence | Reverse primer sequence |
| --- | --- | --- |
| qTaWAK75 | 5- CTTCTTGCCAACACGACCATC -3 | 5- TGTAGCACTGGAACCCGATG -3 |
| qTaWAK76b | 5- CTGCCTCAAATCATCCCTAAGTC -3 | 5- GGATAGTTTTGCCCTCTGACG -3 |
| qActin | 5- AGCCATACTGTGCCAATC -3 | 5- GCAGTGGTGGTGAAGGAGTAA -3 |

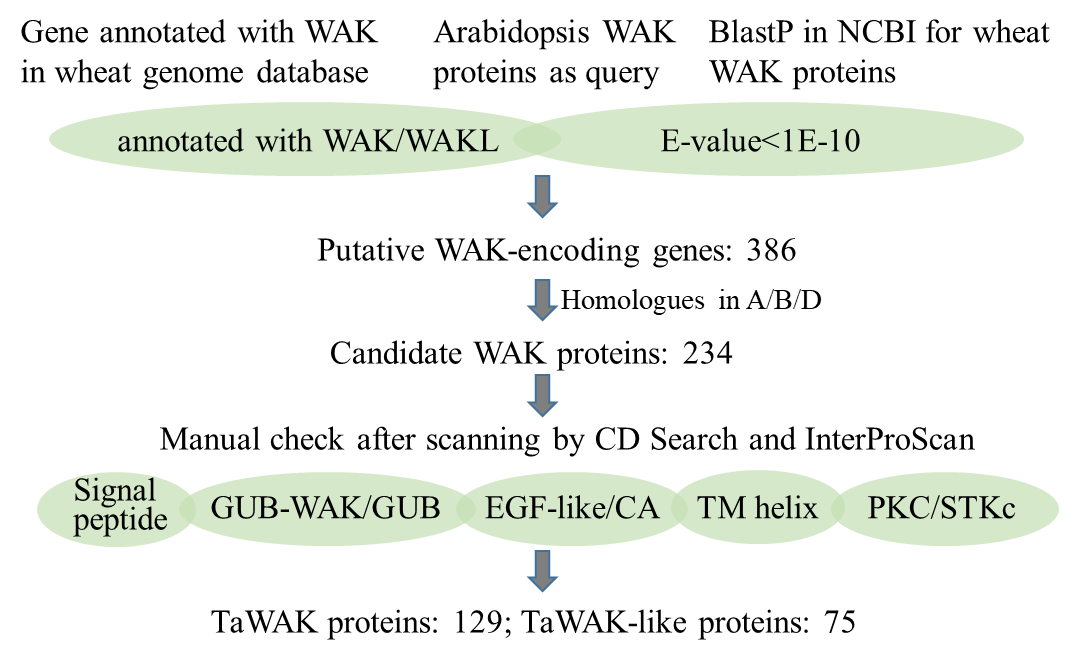

**Figure S1.** Genome-wide identification pipeline of WAKs in wheat genomes. All proteins annotated in IWGSC (RefSeq v1.0) with ‘WAK’ were downloaded, meanwhile five Arabidopsis WAK genes were used as the query sequences for a BLASTp (E-value < 1.0e-50) against NCBI database. The screening results were manually refined using protein domain prediction tools to eliminate non-WAKs after classification of homologues in A, B and D subgenome of wheat.

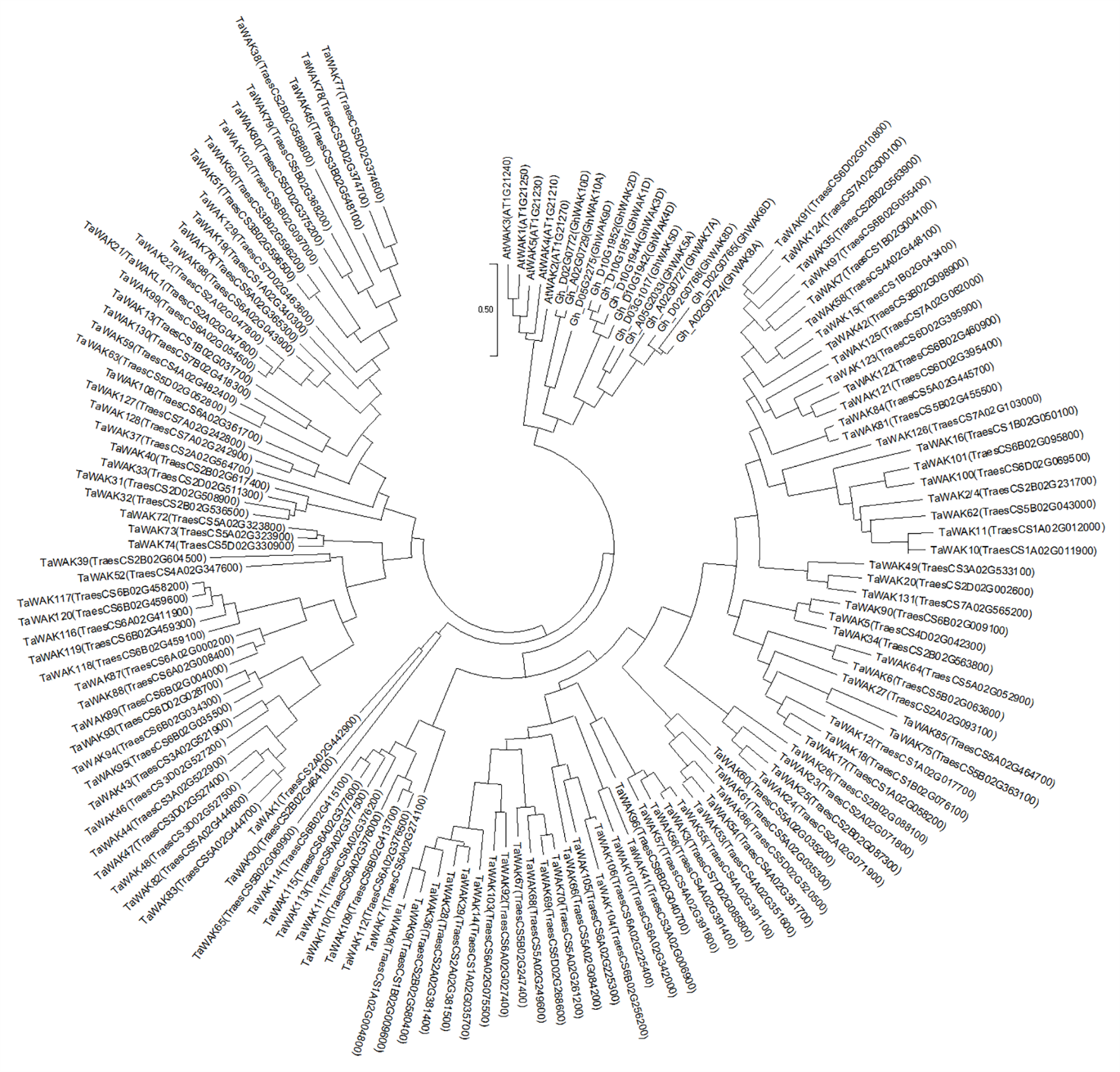

**Figure S2.** Maximum likelihood phylogenetic tree of WAK proteins from wheat, Arabidopsis and cotton using the MEGA software (version 10.0.5). The whole protein sequences were used in the comparison.

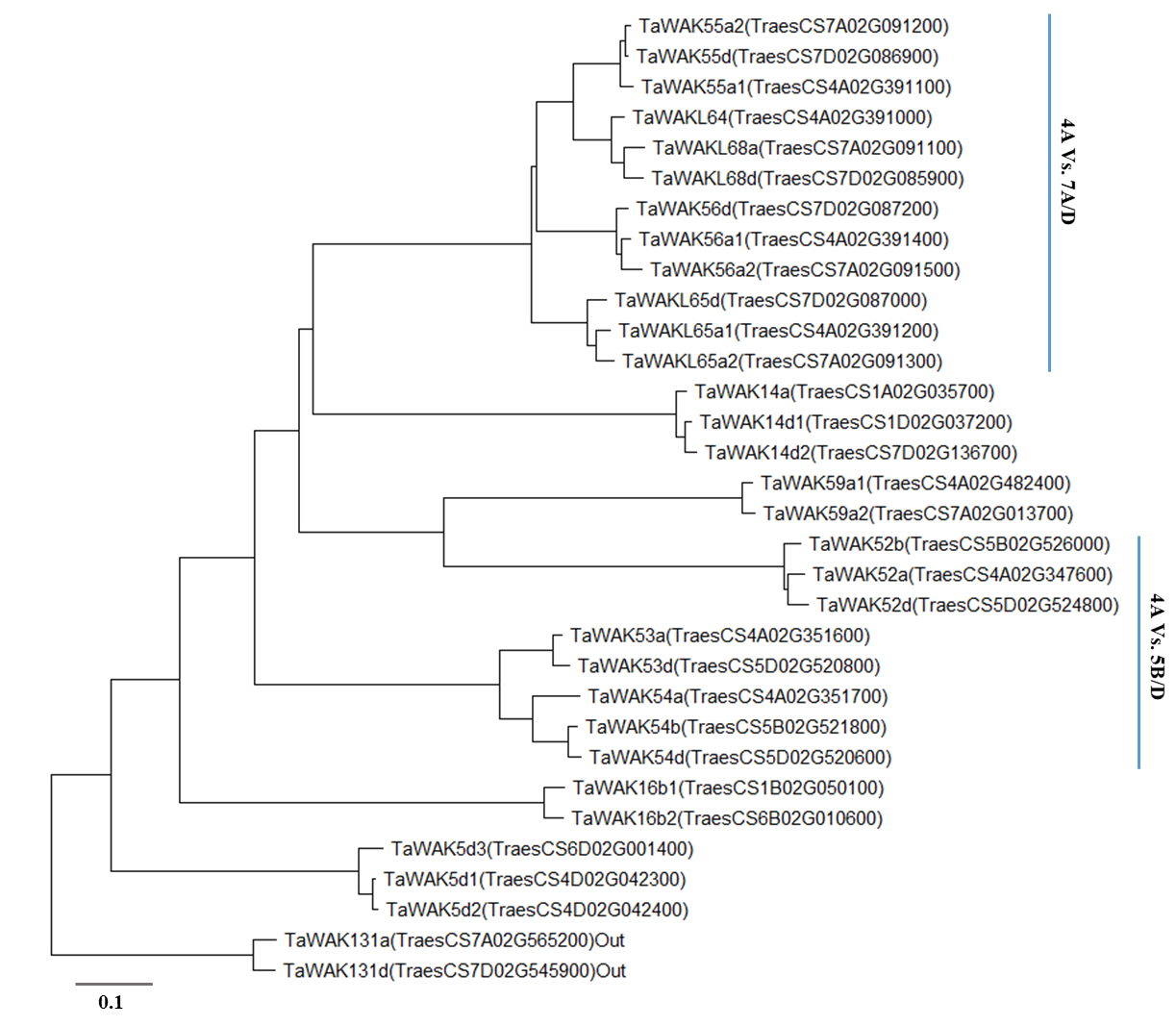

**Figure S3.** Phylogenetic tree of WAK homologues proteins from differently partial homologous group of wheat using the MEGA software (version 10.0.5). The WAK131 protein sequences were used in the comparison as outline protein. The WAK proteins encoding from 4A versus 7A/D or 5B/D were indicated with vertical line.

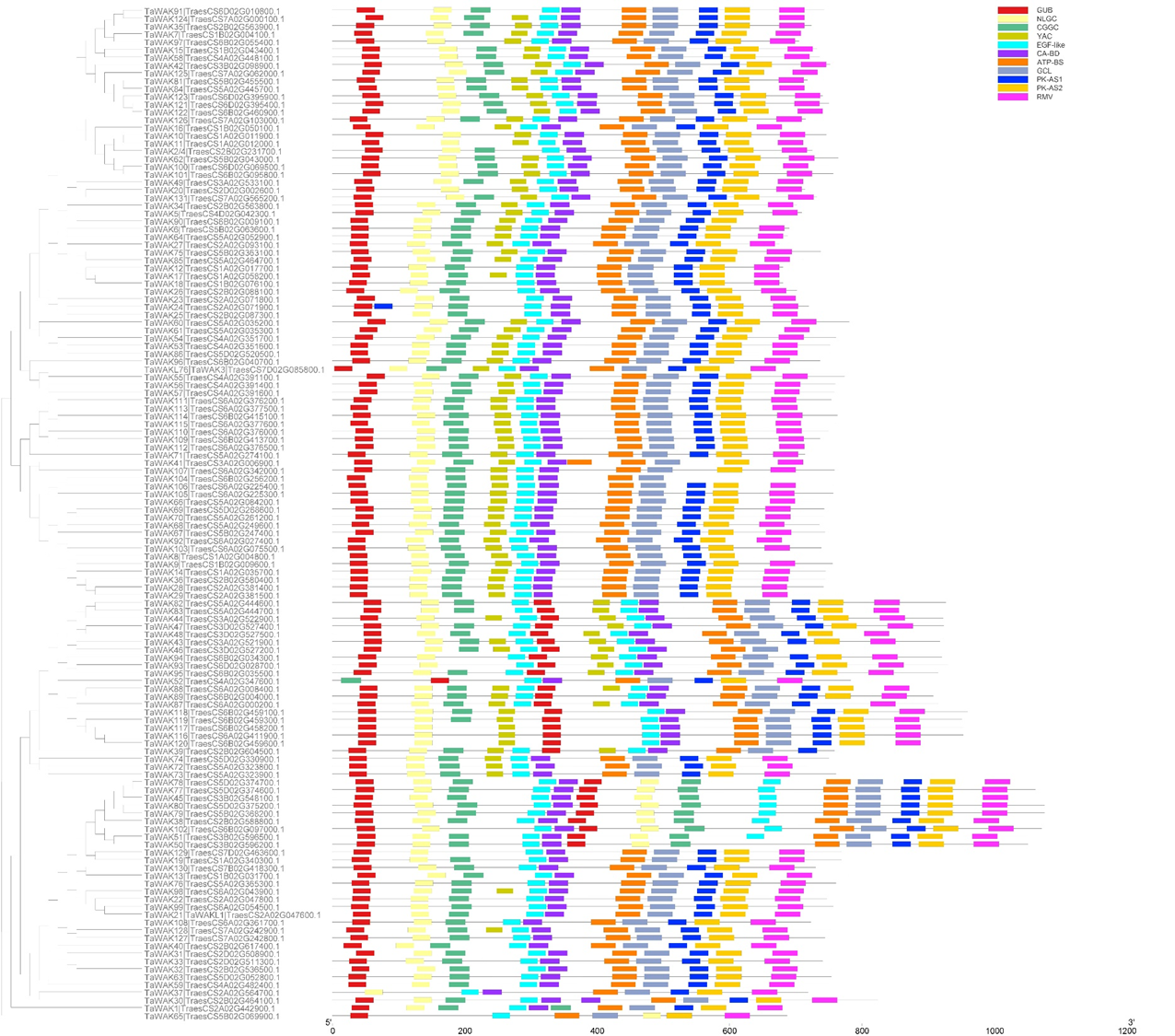

**Figure S4.** Phylogenetic tree, conserved motifs, and sequence structure of 129 TaWAK proteins. The conserved motifs were listed as differential colored box. The sequence structure of TaWAK proteins were represented with the mixed thin line and color boxes, while the scale was given in the bottom of panel.
